# Furin cleavage of the SARS-CoV-2 spike is modulated by O-glycosylation

**DOI:** 10.1101/2021.02.05.429982

**Authors:** Liping Zhang, Matthew Mann, Zulfeqhar Syed, Hayley M. Reynolds, E Tian, Nadine L. Samara, Darryl C. Zeldin, Lawrence A. Tabak, Kelly G. Ten Hagen

## Abstract

The SARS-CoV-2 coronavirus responsible for the global pandemic contains a unique furin cleavage site in the spike protein (S) that increases viral infectivity and syncytia formation. Here, we show that O-glycosylation near the furin cleavage site is mediated by specific members of the GALNT enzyme family and is dependent on the novel proline at position 681 (P681). We further demonstrate that O-glycosylation of S decreases furin cleavage. Finally, we show that GALNT family members capable of glycosylating S are expressed in human respiratory cells that are targets for SARS-CoV-2 infection. Our results suggest that O-glycosylation may influence viral infectivity/tropism by modulating furin cleavage of S and provide mechanistic insight into the potential role of P681 mutations in the recently identified, highly transmissible B.1.1.7 variant.

## Introduction

The novel severe acute respiratory syndrome coronavirus-2 (SARS-CoV-2), which is responsible for the current coronavirus disease 2019 (COVID-19) pandemic, contains a unique insertion of 4 amino acids (PRRA) at the S1/S2 boundary of the spike protein (S) that is not present in SARS-CoV and other related coronaviruses (Fig. 1A). This insertion generates a furin cleavage site that has been shown to increase pseudoviral infectivity (*1–3*) and syncytia formation in cell culture (*4*) (Fig. 1A). Recent structural studies have shown the presence of both prefusion (containing S1 and S2) and postfusion (containing S2 only) S structures on virions independent of target cells (*5*), indicating that the furin site is being used as virions are made in host cells. The authors suggest that the furin-cleaved S2 structures may play a role in immune system evasion, further highlighting the importance of furin cleavage and modifying factors in viral infectivity/tropism. The region around the furin cleavage site contains predicted sites of O-glycosylation, two of which have recently been confirmed in cell culture (*6*) (Fig. 1A). O-glycosylation is known to both positively and negatively influence proteolytic cleavage events in diverse proteins (*7, 8*), suggesting the possibility that furin cleavage of S may be modulated by O-glycosylation.

**Fig. 1.**
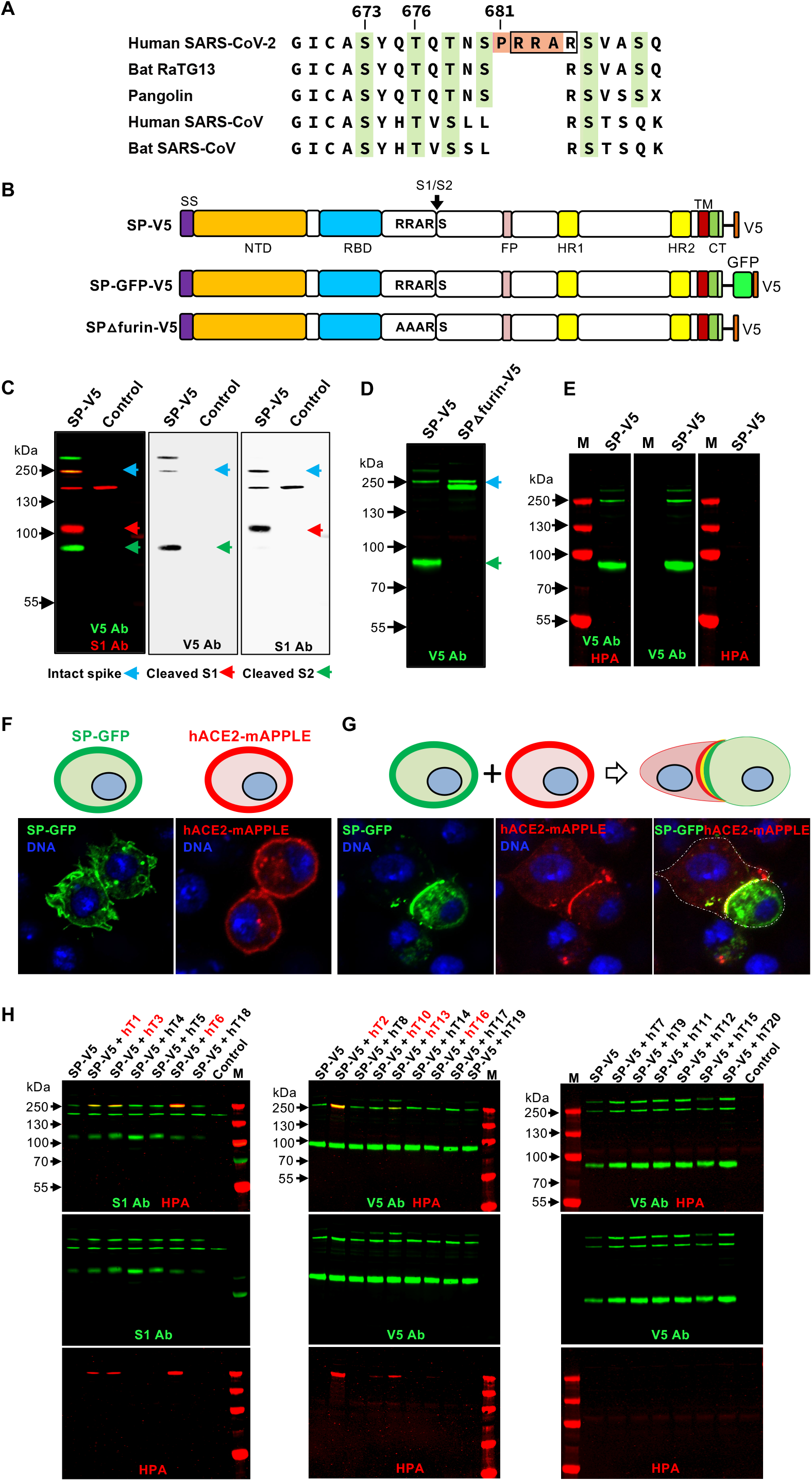
Specific GALNTs O-glycosylate the SARS-CoV-2 S protein. (**A**) Sequence alignment of the S1/S2 region of related betacoronaviruses, highlighting the PRRA insertion unique to SARS-CoV-2 (in orange) and potential sites of O-glycosylation (in green). The furin cleavage site is boxed. Numbering corresponds to the human SARS-CoV-2 sequence. (**B**) Diagram of fulllength WT SARS-CoV-2 S protein (SP) or version with the furin site mutated (RRAR to AAAR; SPΔfurin) linked to either V5 or GFP-V5. The site of furin cleavage at the S1/S2 junction is shown with an arrow. NTD, N-terminal domain; RBD, receptor-binding domain; FP, fusion peptide; HR1, heptad repeat 1; HR2, heptad repeat 2; TM, transmembrane domain; CT, cytoplasmic tail. (**C**) Expression of SP-V5 in *Drosophila* S2R+ cells results in full length S (blue arrow) (detected by the V5 antibody; V5 Ab) along with S1(red arrow) (detected by the S1 antibody; S1 Ab) and S2 (green arrow) (detected by the V5 Ab) cleavage products. (**D**) Mutation of the furin cleavage site results in the presence of only full length S (detected by V5 Ab; green). (**E**) No evidence of O-glycosylation (via HPA staining; red) is seen for S (detected by V5 Ab; green) in this cell background. (**F**) Cells were transfected with either SP-GFP-V5 (green) or the human ACE2 receptor (hACE2-mAPPLE; red) to demonstrate cell surface expression. (**G**) Cells expressing either SP-GFP-V5 or hACE2 were then mixed as shown in the diagram and imaged to reveal that S expressed in these cells maintains the ability to bind to the hACE2 receptor. Nuclear staining is shown in blue. (**H**) SP-V5 was cotransfected with each member of the human *GALNT* family (hT1-20) and O-glycosylation was assessed by staining with the O-glycan-specific lectin HPA (red). S was detected via staining with the S1 Ab (red) or V5 Ab (green). GALNT family members that glycosylate S are shown in red at the top of the panel. Control is the empty vector. M, markers. Marker size is shown in kDa on the side of each panel.

## Results and Discussion

To investigate the role of O-glycosylation in modulating furin cleavage of S, we expressed GFP- or V5-labeled full-length S with or without the novel furin cleavage site (Fig. 1B) in a cell culture background where no O-glycosylation was detected and furin cleavage was active. As shown in Fig. 1C, S expressed in *Drosophila* S2R+ cells results in full-length S as well as S1 and S2 cleavage products that are the result of furin cleavage; mutation of the furin site (RRAR to AAAR; SPΔfurin-V5) abrogated cleavage, resulting in the presence of only intact S (Fig. 1D). Likewise, treatment with furin inhibitors decreased cleavage (Fig. S1A). S expressed in these cells displayed no reactivity to the O-glycan-specific lectins *Helix pomatia* (HPA) or *Peanut agglutinin* (PNA) (Fig. 1E and Fig. S1B), confirming no detectable background O-glycosylation. Additionally, S expressed in these cells retained functional binding ability, as evidenced by binding to cells expressing the receptor hACE2 (Fig. 1F-G). We therefore used this cell background to screen for members of the human UDP-GalNAc:polypeptide *N*-acetylgalactosaminyltransferase (GALNT) enzyme family (*9*) that are capable of glycosylating S. Interestingly, only certain GALNTs were effective at glycosylating S (Fig. 1H). S glycosylation (as detected by HPA reactivity) was detected upon coexpression with GALNT1, 2, 3, 6, 10, 13 or 16 (Fig. 1H). No glycosylation was seen upon coexpression with GALNT4, 5, 7, 8, 9, 11, 12, 14, 15, 17, 18, 19 or 20 (Fig. 1H).

We next examined which of the identified GALNTs are capable of glycosylating the furin proximal region of S. GALNT1, 2, 3, 6, 13 and 16 were expressed, partially purified and tested for GalNAc transfer (as described previously; (*10*)) to peptides within the furin proximal region of SARS-CoV-2 S. Enzyme assays demonstrated that only GALNT1 showed detectable transferase activity above background against this region of S (Fig. 2A-B and Fig. S2A-F). Additionally, examining the activity of GALNT1 against peptides where each potential site of O-glycosylation was eliminated revealed a dramatic loss of activity against a peptide containing mutations at T678, suggesting that this is a preferred site of GalNAc addition by GALNT1 (Fig. 2A-B and Fig. S2A). T678 is one of the sites previously identified as being O-glycosylated in human cells by mass spectroscopy (*6*). Likewise, a decrease in GalNAc transfer was seen with a peptide containing mutations at T676, suggesting this is also a site of addition for GALNT1 (Fig. 2A-B and Fig. S2A). Interestingly, complete abrogation of all GALNT1-mediated glycosylation was seen upon mutating the unique proline at position 681 (P681), indicating that GALNT1 activity in this region is dependent on this proline. No glycosylation of the comparable region of SARS-CoV-1 (that lacks the novel PRRA insertion) by GALNT1 was observed. Taken together, these results suggest GALNT1 glycosylates specific residues within the furin proximal region of the SARS-CoV-2 S and that glycosylation by GALNT1 is dependent on the presence of P681.

**Fig. 2.**
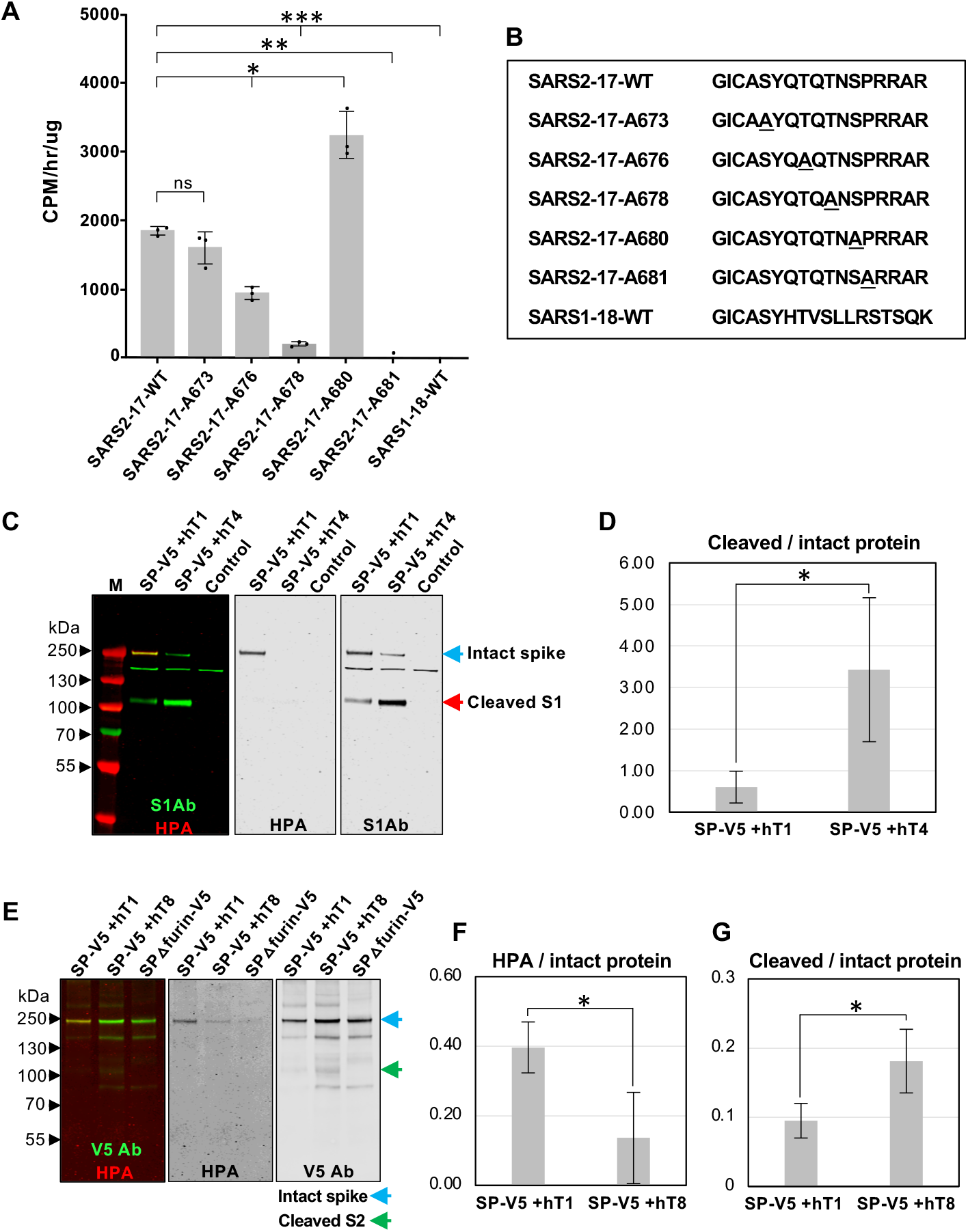
O-glycosylation of SARS-CoV-2 S decreases furin cleavage. (**A**) Enzyme assays showing GALNT1 activity on specific residues in the region of S proximal to the furin cleavage site and the S1/S2 border. GALNT1 glycosylates this region of SARS-CoV-2 (SARS2-17-WT) but not of SARS-CoV-1 (SARS1-18-WT). GALNT1 activity is dependent on the unique proline at position 681 (P681). (**B**) Peptides used in enzyme assays. Mutated residues are underlined. (**C**) Coexpression of S with GALNT1 (hT1) (which glycosylates S) results in decreased furin cleavage relative to coexpression with GALNT4 (hT4) (which does not glycosylate S) in *Drosophila* S2R+ cells. The S1Ab was used to assess the ratio of cleaved to uncleaved S (denoted by arrows). O-glycosylation is seen (HPA staining) only on the intact S coexpressed with GALNT1. (**D**) Average ratios of cleaved/intact S cotransfected with either GALNT1 or GALNT4 from 3 independent experiments in *Drosophila* S2R+ cells are shown. (**E**) Coexpression of S with GALNT1 in Vero E6 cells results in increased O-glycosylation of S (**F**), as detected by HPA and decreased furin cleavage (**G**) relative to coexpression with GALNT8 (hT8) (which does not glycosylate S). The V5 Ab was used to assess the ratio of cleaved to uncleaved S (denoted by arrows). M, markers. Marker size is shown in kDa on the side of each panel. Error bars are SD. \**P*<0.05;\*\**P*<0.01;\*\*\**P*<0.001.

Our cell culture experiments revealed that O-glycosylation was only present on the full-length S, and never seen on either the S1 or S2 proteolytic fragments (Fig. 1H), suggesting that these events are mutually exclusive. To investigate whether O-glycosylation may interfere with furin cleavage of S, we coexpressed S with various GALNTs and assessed the ratio of cleaved to intact S. S was cotransfected with either GALNT1 (which glycosylates S) or GALNT4 (which does not glycosylate S) and ratios of S were quantitated via western blots. As shown in Fig. 2C-D and Fig. S3, coexpression of S with GALNT1 resulted in O-glycosylation of S and dramatically reduced the degree of furin cleavage. To examine whether O-glycosylation modulates furin cleavage of S in mammalian cells, we performed the same experiments in Vero E6 cells, comparing the effects of GALNT1 to GALNT8 (which does not glycosylate S). As shown in Fig. 2E-G and Fig. S4, coexpression of S with GALNT1 resulted in increased O-glycosylation of intact S and decreased furin cleavage, relative to coexpression with GALNT8. Taken together, our results suggest that O-glycosylation of S by specific members of the GALNT family can modulate furin cleavage and the ratio of cleaved to intact S produced in mammalian cells.

Given the potential role of endogenously expressed GALNTs to alter the processing of S in cells, we next set out to define the repertoire of *GALNTs* expressed in cells of the respiratory tract likely to be infected by SARS-CoV-2. We mined the single-cell RNA sequencing databases from cells of the lower respiratory tract of healthy controls from Travaglini et al., 2020 (*11*). As shown in Fig. 3, *GALNT1, 2, 3, 6, 7, 11, 12* and *18* are the predominant family members expressed. In particular, abundant expression of *GALNT1* is notable across many cell types that also express ACE2, and are thus likely targets for SARS-CoV-2 infection (Fig. 3A-B). We next examined expression levels of *GALNTs* in the upper respiratory cells of healthy controls from the dataset of Chua et al., 2020 (*12*) (Fig. 3C-D). In cells expressing the highest levels of *ACE2* (ionocytes), *GALNT1* is the most abundantly expressed family member (Fig. 3C-D).

**Fig. 3.**
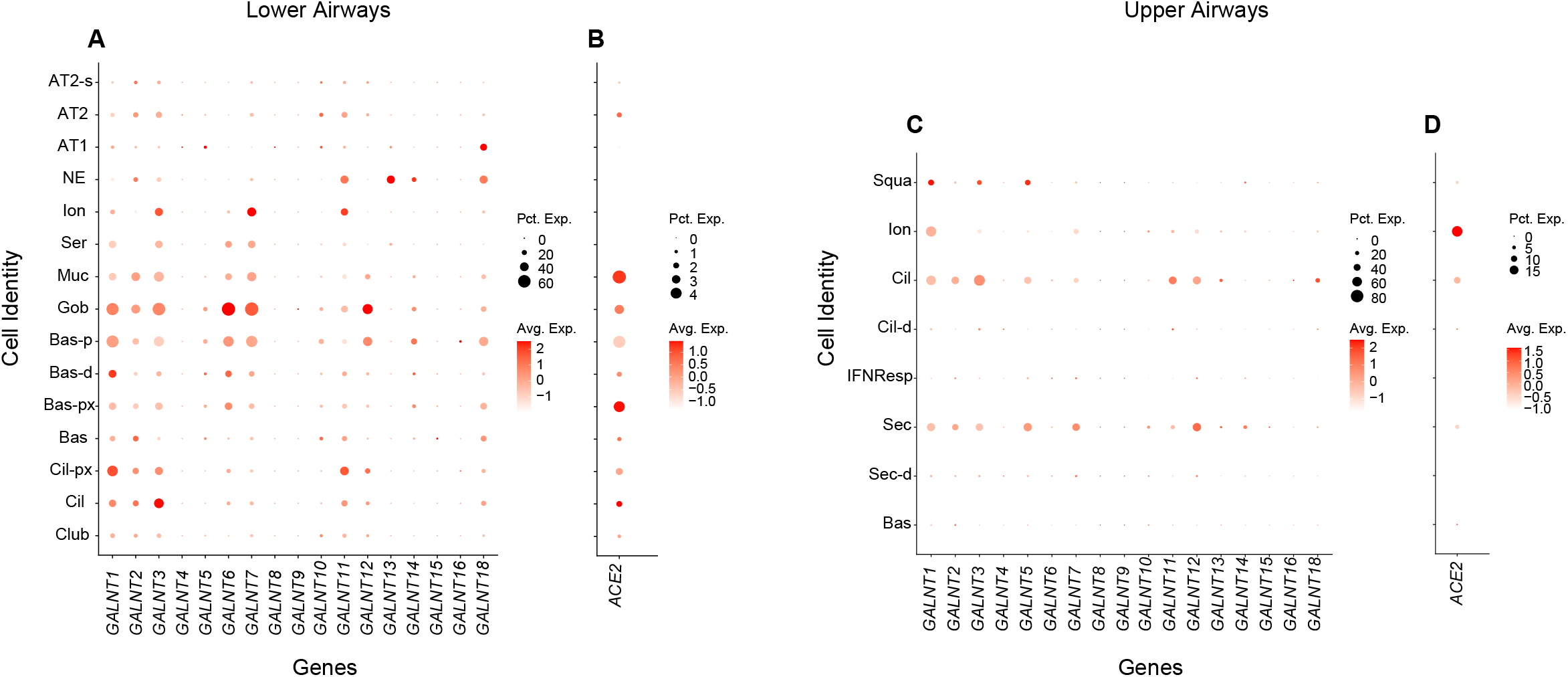
*GALNT* expression in cells of the human respiratory tract and human cell lines. Dot plots showing expression of *GALNT* family members (**A**) and *ACE2* (**B**) in cells of the lower respiratory tract from the cell atlas of a healthy control (*11*). Dot size represents the percentage of cells expressing each gene and color intensity denotes degree of expression with each group. Cell types shown are: AT1, alveolar epithelial type-1; AT2, alveolar epithelial type-2; AT2-s, AT2-signalling; NE, neuroendocrine; Ion, ionocyte; Ser, serous; Muc, mucous; Gob, goblet; Bas-p, proliferating basal; Bas-d, differentiating basal; Bas-px, proximal basal; Bas, basal; Cil-px, proximal ciliated; Cil, ciliated; Club. Dot plots showing expression of *GALNT* family members (**C)** and *ACE2* (**D**) in cells of the upper respiratory tract from the cell atlas of a healthy control (*12*). Cell types shown are: Squa, squamous; Ion, ionocyte; Cil, ciliated; Cil-d, differentiating ciliated; IFNResp, IFNG responsive; Sec, secretory; Sec-d, differentiating secretory; Bas, basal.

Our results identify O-glycosylation as a modulator of furin cleavage of the SARS-CoV-2 S protein. Given the role of the furin cleavage in increased infectivity and syncytia formation, our results suggest that O-glycosylation may alter viral tropism and/or infectivity via influencing the degree to which S is processed by furin, raising the possibility that differences in *GALNT* expression from individual to individual could influence disease severity. In particular, we identify one O-glycosyltransferase, GALNT1, that is expressed within ACE2+ cells of the upper and lower respiratory tracts, and specifically glycosylates the furin proximal region within S. Additionally, the activity of GALNT1 within the furin proximal region is dependent on the unique P681 present in SARS-CoV-2, which is supported by previous studies indicating that GALNT1 has a strong preference for substrates with vicinal prolines (*13*). Given that the recently identified B.1.1.7 variant contains mutations in P681, our results suggest that the effect of the loss of that proline on glycosylation of S may be a contributing factor to its increased infectivity.

## Materials and Methods

### Cloning of SARS-CoV-2 *Spike (S*), human *ACE2* and *GALNTs*

The codon optimized cDNA of full-length *spike* (Genscript) was amplified by PCR and digested by HindIII and NotI, then cloned into pIB/V5-His vector (Invitrogen), fused with V5 epitope at the C-terminus. To make mutations to remove the furin cleavage site (spikeΔfurin), two arginines at position 682 and 683 were changed to alanines using the QuickChange II XL site-directed mutagenesis kit (Agilent Technologies) and the plasmid pIB-spike-V5 as a template. Mutagenic primers were designed using the QuickChange Primer Design Program available online. The sense primer sequence is 5′-cgagcacggcgagctgagttagtctggg-3′ and anti-sense primer sequence is 5′-cccagactaactcagctcgccgtgctcg-3′. Reactions and transformation were performed according to the manufacturer’s instructions. DNA sequencing was performed to verify the mutations. The cDNA of *hACE2* (Genscript) was amplified by PCR and digested by KpnI and NotI, then cloned into pIB/V5-His vector, fused with mAPPLE/V5 epitope at the C-terminus. The cDNAs of *GALNTs* 1-20 (Genscript) were amplified by PCR and digested by BamHI and NotI (*GALNT1, 2, 3, 4, 9, 13, 14, 17, 18, 19, 20*) or HindIII and NotI (*GALNT5, 6, 8, 10, 11, 15, 16*) or HindIII and XhoI (*GALNT12*), or SacI and NotI (*GALNT7*), then cloned into the pIB vector, fused with FLAG epitope at the C-terminus.

pcDNA3.1-spike and pcDNA-3.1-spikeΔfurin (Genscript) were fused with V5 epitope at the C-terminus to make pcDNA3.1-spike-V5 and pcDNA-3.1-spikeΔfurin-V5.

### Expression of Spike and GALNTs in *Drosophila* S2R+ cells/Vero E6 cells and western blotting

The plasmids pIB-spike-V5 with or without pIB-GALNTs-FLAG were transfected to S2R+ cells (DGRC) using Effectene transfection reagent (Qiagen) according to the instructions. For expression of Spike in Vero E6 cells (ATCC), the cells were transfected with pcDNA3.1-spike-V5 with or without pcDNA3.1-GALNTs-FLAG (Genscript) using Lipofectamine 3000 reagent (Invitrogen) according to the instructions.

For western blotting, cells were lysed with RIPA buffer (Sigma) containing 1X Halt Protease Inhibitor (Thermo Scientific) 3-4 days after transfection. Protein extracts were incubated with agarose-immobolized V5 antibody (Bethyl) overnight at 4°C. Then beads were collected and washed with PBS four times and protein samples were mixed with LDS sample buffer (Invitrogen) containing BME. Proteins were analyzed by NuPAGE 4–12% Bis-Tris gels and transferred onto nitrocellulose membranes. The membranes were blocked with Odyssey Blocking Buffer (PBS-based) (Li-COR), then incubated with anti-V5 antibody (dilution 1:1000, Invitrogen) or anti-S1 antibody (dilution 1:1000, GeneTex) overnight at 4 °C. After washing with PBS containing 0.1% Tween-20 (PBST), the membrane was incubated with IRDye 680LT-conjugated anti-mouse IgG (1:5000, Li-COR) or 680LT-conjugated anti-rabbit IgG (1:5000, Li-COR) and IRDye 800CW (Li-COR) labeled HPA (Helix pomatia lectin, Sigma) or PNA (peanut agglutinin, Sigma) for 1hr at room temperature. After washing with PBST and PBS, the membranes were scanned by a Li-COR Odyssey Infrared Imaging System. The protein band intensity was measured using ImageJ to calculate the ratio of cleaved protein/ intact protein and the ratio of HPA/ intact protein.

### Furin inhibitor treatment in cell culture

24 hrs after transfection, the furin inhibitor decanoyl-RVKR-CMK (Tocris) was added to the medium at final concentrations of 20uM, 50uM or 100uM. Cells were collected 72 hrs after transfection and lysed with RIPA buffer (Sigma) containing 1X Halt Protease Inhibitor (Thermo Scientific). The cell lysates were used for immunoprecipitation and western blotting experiments.

### Cell imaging

Cells were fixed with 4% PFA, washed with PBST and mounted with Vectashield Mouting Medium with DAPI (Vector Laboratories) 2 days after transfection. The cells were imaged on Nikon A1R confocal microscope. Images were processed using ImageJ.

### Enzyme assays

Expression of recombinant GALNTs was performed either using *Pichia pastoris* or COS7 cells as described previously and referenced in the main text. Peptide substrates were synthesized by Peptide 2.0. Assays were performed as described previously and referenced in the main text. Reactions were run for 30 min at 37°C in 25 μl final volumes consisting of the following: 25 mM acceptor substrate, 7.3 μM UDP-[^14^C]-GalNAc (54.7 mCi/mmol; 0.02mCi/ml), 2 mM cold UDP-GalNAc, 500 mM MnCl_2_, 40 mM cacodylate (pH 6.5), 40 mM 2-mercaptoethanol and 0.1 % Triton X-100. Reactions were then quenched with 30 mM EDTA. Glycosylated products were separated from unincorporated UDP-[^14^C]-GalNAc by anion exchange chromatography using AG1-X8 resin columns (Bio-Rad #1401454), and product incorporation was determined by liquid scintillation counting (Beckman Coulter LS6500). Reactions without acceptor peptide were also used to generate background values that were subtracted from each experimental value. Assays for each peptide substrate were run in triplicate and repeated three times. Experimental values for each substrate were then averaged and standard deviations were calculated. Enzyme activity (initial rate) is expressed as CPM/hr/μM or CPM/hr.

### Quantitative real-time PCR

DNase-free RNA was isolated by using PureLink RNA Mini Isolation Kit (Ambion). cDNA synthesis was performed by using the iScript cDNA Synthesis Kit (Bio-Rad). Human primers were designed by using Beacon Designer software (Bio-Rad). Quantitative PCR was performed on a CFX96 real-time PCR thermocycler (Bio-Rad) using the SYBR-Green PCR Master Mix (Bio-Rad). Quantitative PCR was performed in triplicate. Gene expression levels were normalized to human *29S* rRNA and plotted as gene expression level and relative expression levels to a housekeeping gene. Values represent mean value.

### scRNASeq Analysis

Published datasets for upper and lower airways were retrieved as R data format (.rds) and the encapsulated Seurat (v.3.2.3) object was updated to Seurat v3 object using the Seurat UpdateSeuratObject() function. We subset the data for the Epithelial cell compartment from the upper and lower airways atlas and the expression of *GALNTs* and *ACE2* is illustrated as dot plots using the R ggplot2 (v.3.3.2) library. Individual dot plots for upper and lower airways were assembled in Adobe Illustrator (v.24.3) to generate a final figure. The datasets are available for download from their respective published sources referenced in the main text.

## Acknowledgments

We would like to thank our colleagues for many helpful discussions.

## Author Contributions

L. Z., E. T., H. R., Z. S., N. L. S., D. Z., L A. T., and K.G.T.H. designed and planned the research. L. Z., E. T., H. R., M. M., N. M. and Z. S. performed experiments, with input from L. A. T. and K.G.T.H. L. Z., E. T., H. R., Z. S., M. M., N. L. S., L.A.T. and K. G. T. H. analyzed and discussed the data. K.G.T.H. wrote the paper with input from L. Z., E. T., H. R., Z. S., M. M., N. L. S., N. M., D. C. Z. and L.A.T.

## Data and materials availability

All data are available in the main text or the supplementary materials.

## References and Notes

1. J. L. Daly et al., Neuropilin-1 is a host factor for SARS-CoV-2 infection. Science 370, 861–865 (2020).

2. L. Cantuti-Castelvetri et al., Neuropilin-1 facilitates SARS-CoV-2 cell entry and infectivity. Science 370, 856–860 (2020).

3. J. Shang et al., Cell entry mechanisms of SARS-CoV-2. Proc Natl Acad Sci U S A 117, 11727–11734 (2020).

4. M. Hoffmann, H. Kleine-Weber, S. Pohlmann, A Multibasic Cleavage Site in the Spike Protein of SARS-CoV-2 Is Essential for Infection of Human Lung Cells. Mol Cell 78, 779–784 e775 (2020).

5. Y. Cai et al., Distinct conformational states of SARS-CoV-2 spike protein. Science 369, 1586–1592 (2020).

6. C. Gao et al., SARS-CoV-2 Spike Protein Interacts with Multiple Innate Immune Receptors. bioRxiv, (2020).

7. L. Zhang et al., O-glycosylation regulates polarized secretion by modulating Tango1 stability. Proc Natl Acad Sci U S A 111, 7296–7301 (2014).

8. K. T. Schjoldager et al., A systematic study of site-specific GalNAc-type O-glycosylation modulating proprotein convertase processing. J Biol Chem 286, 40122–40132 (2011).

9. D. T. Tran, K. G. Ten Hagen, Mucin-type O-glycosylation during development. J Biol Chem 288, 6921–6929 (2013).

10. S. Ji et al., A molecular switch orchestrates enzyme specificity and secretory granule morphology. Nat Commun 9, 3508 (2018).

11. K. J. Travaglini et al., A molecular cell atlas of the human lung from single-cell RNA sequencing. Nature 587, 619–625 (2020).

12. R. L. Chua et al., COVID-19 severity correlates with airway epithelium-immune cell interactions identified by single-cell analysis. Nat Biotechnol 38, 970–979 (2020).

13. T. A. Gerken et al., Emerging paradigms for the initiation of mucin-type protein O-glycosylation by the polypeptide GalNAc transferase family of glycosyltransferases. J Biol Chem 286, 14493–14507 (2011).

