## Supplementary Figures for "Furin cleavage of the SARS-CoV-2 spike is modulated by O-glycosylation"

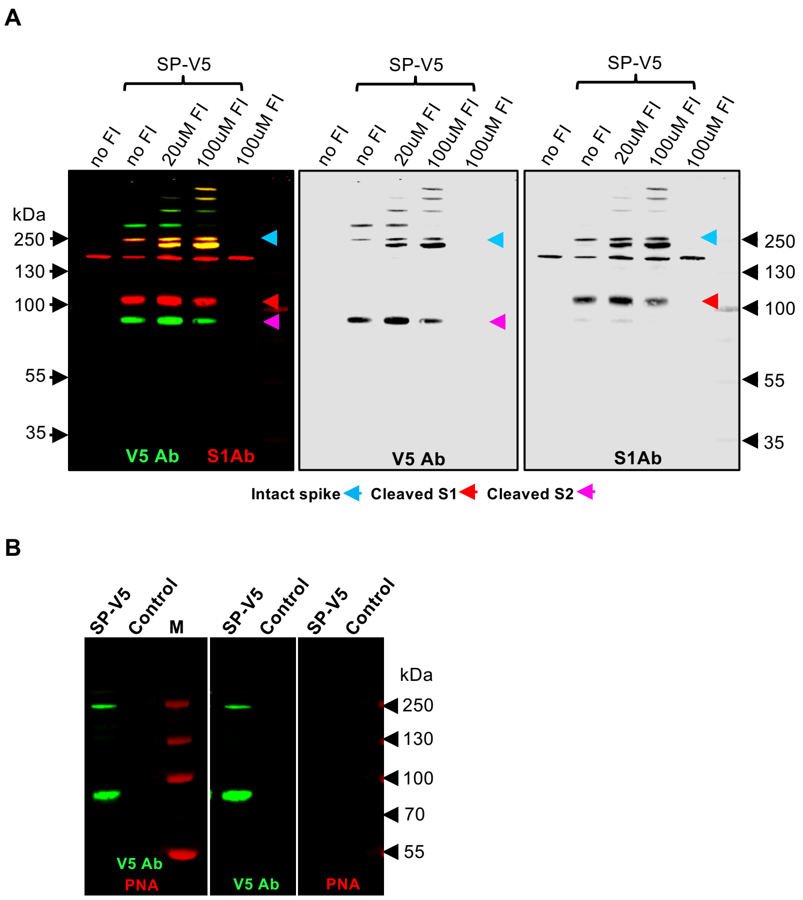


**Fig. S1. Furin cleavage of S expressed in *Drosophila* S2R+ cells.** (**A**) Decreased cleavage of S expressed in *Drosophila* S2R+ cells is seen upon incubation with increased concentrations of a furin inhibitor (FI). V5 Ab was used to detect intact S (blue arrow) and the cleaved S2 fragment (pink arrow). The S1 Ab was used to detect the cleaved S1 fragment (red arrow). (**B**). S expressed in *Drosophila* S2R+ cells (as detected by V5 Ab) is not PNA reactive (red). Size markers (kDa) are shown to the left and right of each blot.

A

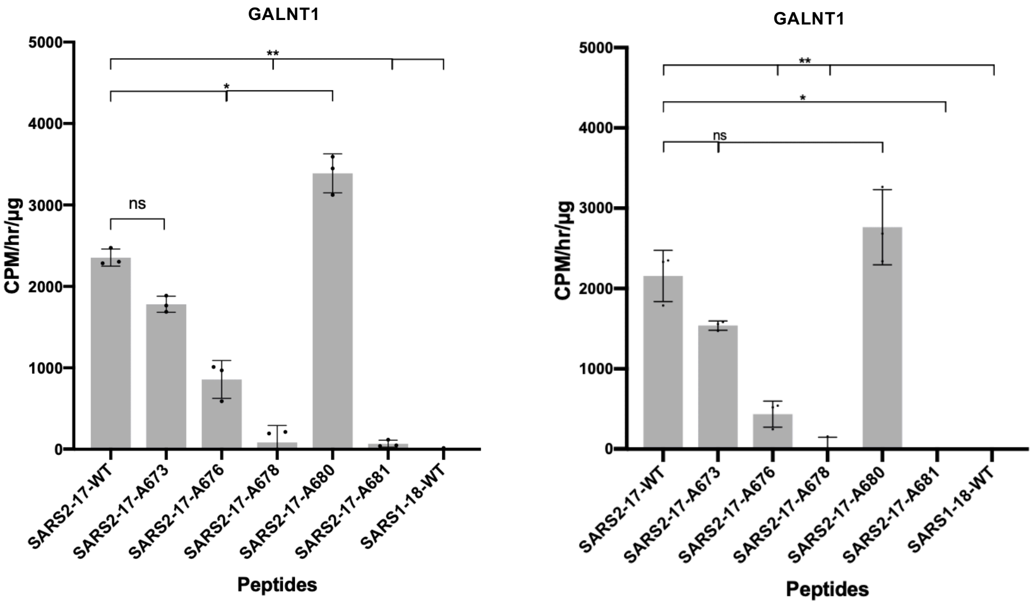
GALNT1
CPM/hr/µg
5000
4000
3000
2000
1000
0
ns
*
**
SARS2-17-WT
SARS2-17-A673
SARS2-17-A676
SARS2-17-A678
SARS2-17-A680
SARS2-17-A681
SARS1-18-WT
Peptides

B

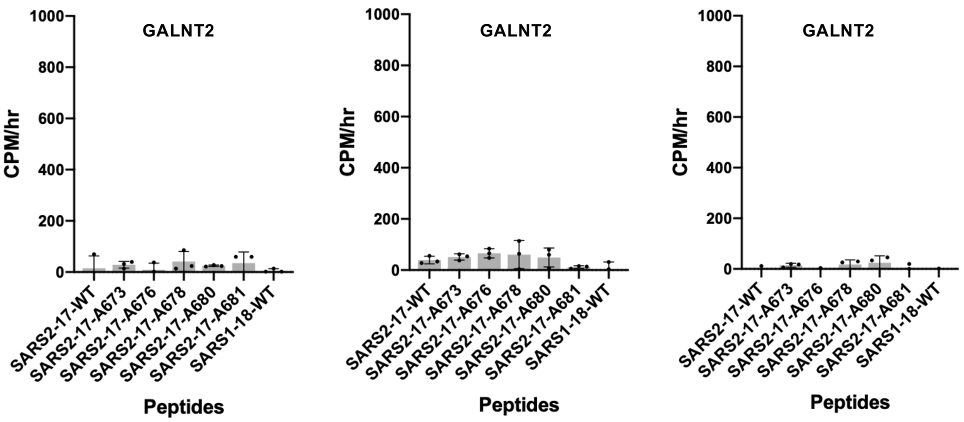
GALNT2
CPM/hr
1000
800
600
400
200
0
SARS2-17-WT
SARS2-17-A673
SARS2-17-A676
SARS2-17-A678
SARS2-17-A680
SARS2-17-A681
SARS1-18-WT
Peptides

C

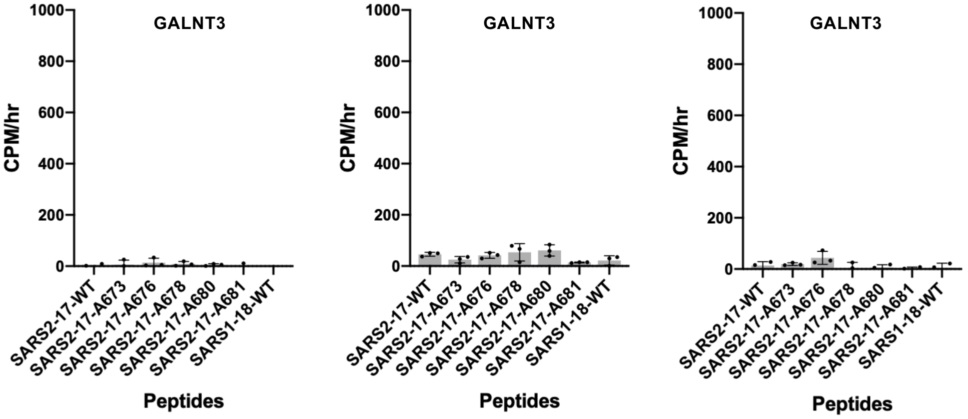
GALNT3
CPM/hr
1000
800
600
400
200
0
SARS2-17-WT
SARS2-17-A673
SARS2-17-A676
SARS2-17-A678
SARS2-17-A680
SARS2-17-A681
SARS1-18-WT
Peptides

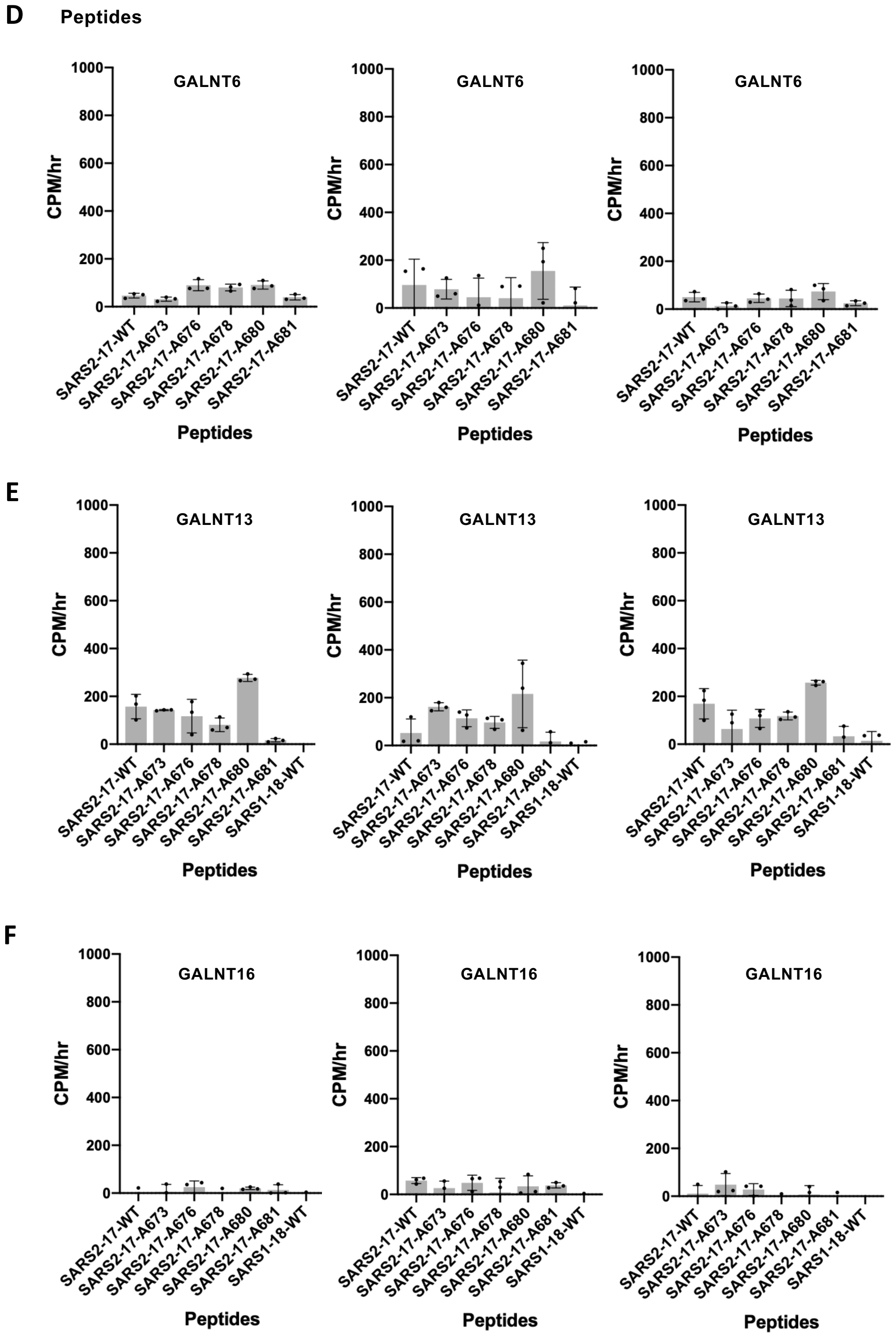


**Fig. S2. O-glycosylation of SARS-CoV-2 S by GALNT1.** Enzyme assays testing for O-glycosylation of peptides and mutated variants from the furin proximal region of the SARS-CoV-2 S and the related region of SARS-CoV-1 S using GALNT1 (**A**), GALNT2 (**B**), GALNT3

(**C**), GALNT6 (**D**), GALNT13 (**E**), and GALNT16 (**F**). GALNT1 activity is dependent on the unique proline at position 681 (P681). Peptide sequences are as shown in the main text. Each data point represents an individual assay. Error bars are SD.*$P<0.05$;**$P<0.01$.

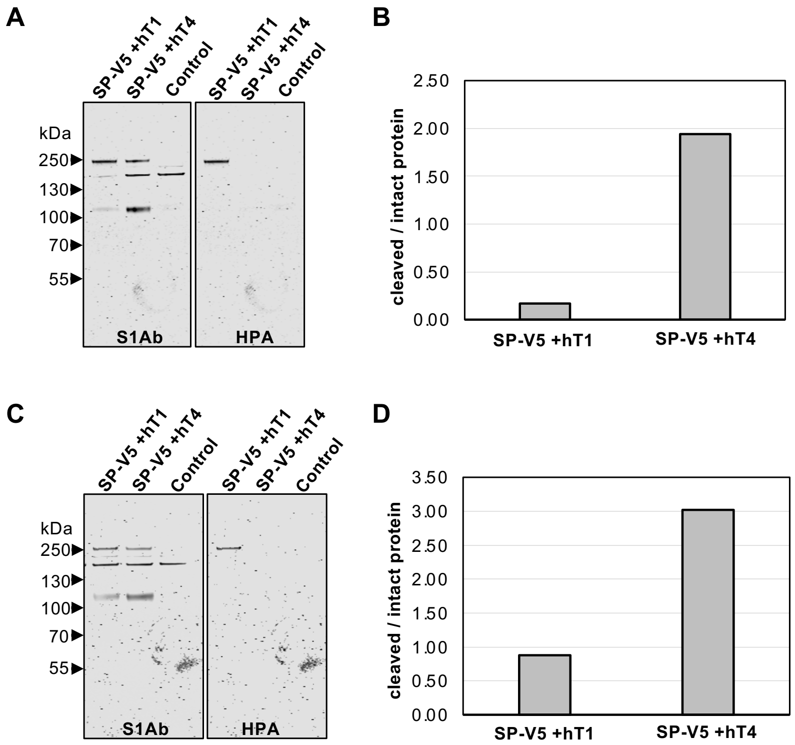


**Fig. S3. O-glycosylation of SARS-CoV-2 S decreases furin cleavage in *Drosophila* S2R+ cells.** Repeats of experiments coexpressing S with GALNT1 (hT1) (which glycosylates S) or GALNT4 (hT4) (which does not glycosylate S) in *Drosophila* S2R+ cells. S1Ab was used to detect cleaved and uncleaved S via westerns (**A** and **C**). O-glycosylation is seen (HPA staining) only on the intact S coexpressed with GALNT1. Ratios of cleaved to uncleaved S were quantitated (**B** and **D**).

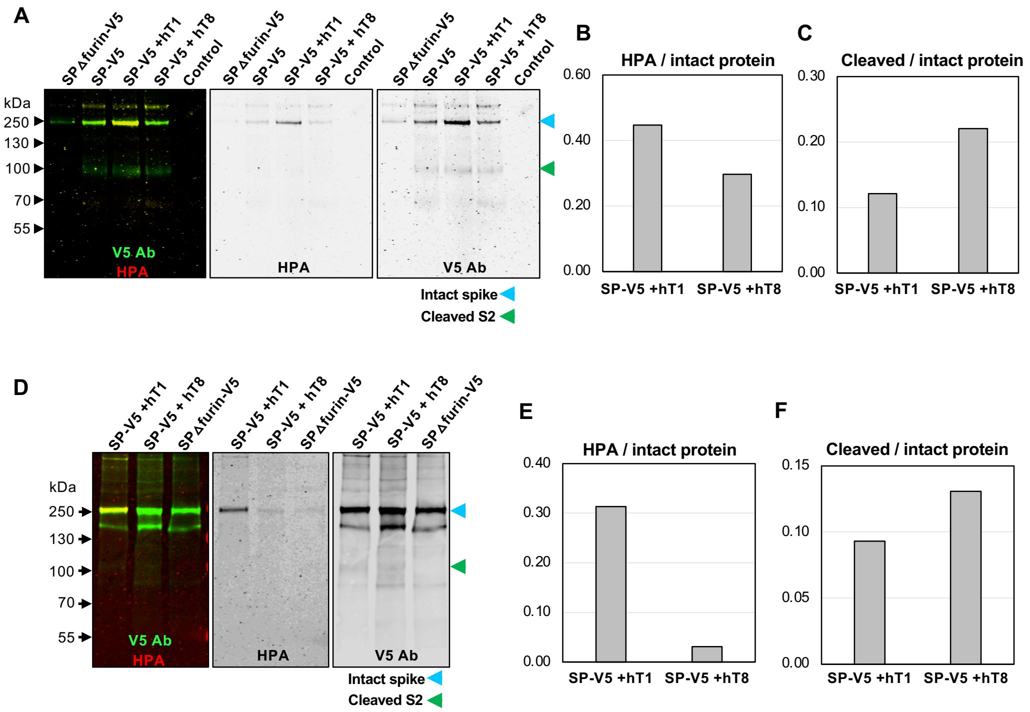


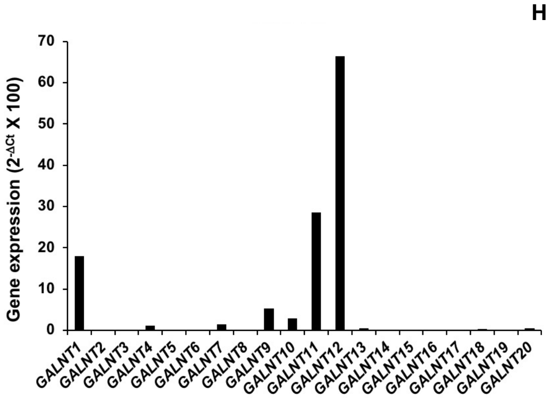


H

| Gene | CT | SD |
|---|---|---|
| *hS29* | 19.14 | 0.12 |
| *GALNT1* | 24.94 | 0.07 |
| *GALNT2* | ND | NA |
| *GALNT3* | 33.64 | 0.52 |
| *GALNT4* | 29.02 | 0.20 |
| *GALNT5* | 36.36 | 0.62 |
| *GALNT6* | 36.22 | 1.09 |
| *GALNT7* | 28.58 | 0.07 |
| *GALNT8* | ND | NA |
| *GALNT9* | 26.69 | 0.43 |
| *GALNT10* | 27.60 | 0.08 |
| *GALNT11* | 24.27 | 0.03 |
| *GALNT12* | 23.05 | 0.04 |
| *GALNT13* | 30.34 | 0.44 |
| *GALNT14* | ND | NA |
| *GALNT15* | ND | NA |
| *GALNT16* | 33.35 | 0.70 |
| *GALNT17* | 34.12 | NA |
| *GALNT18* | 30.80 | 0.16 |
| *GALNT19* | 37.65 | 0.78 |
| *GALNT20* | 30.28 | 0.30 |

**Fig. S4. O-glycosylation of SARS-CoV-2 S decreases furin cleavage in Vero E6 cells.** Repeats of experiments coexpressing S with GALNT1 (hT1) (which glycosylates S) or GALNT8 (hT8) (which does not glycosylate S) Vero E6 cells. V5 Ab was used to detect cleaved and uncleaved S via westerns (**A** and **D**). O-glycosylation (HPA staining) is increased on intact S coexpressed with GALNT1 relative to GALNT8 (**B** and **E**). Ratios of cleaved to uncleaved S from each experiment were quantitated (**C** and **F**). Endogenous expression of *GALNT* family members in Vero E6 cells was assessed by QPCR (**G**). Cycle threshold (CT) values (**F**) are shown. Error bars are SD. ND, not detected. NA, not available.

**Table S1.**
**PCR primer sequences**

| Gene | Sense primer sequence | Anti-sense primer sequence |
|---|---|---|
| *GALNT1* | GCAGGCAGTGCTATTTCTTGTG | CTGCTTCCTTTCAAATTTCCTGGG |
| *GALNT2* | CCTCTCAGCCTTCCTATCATCC | CGCAACCGCAGTAAGTCACT |
| *GALNT3* | CTCGGTTCTGCCTCTCCA | ACAGTTGCGGCTCAGTAGA |
| *GALNT4* | GCGGTGAGGTGGACTTGG | GGAGGTCTGAGAGCCTTCTTGA |
| *GALNT5* | CCACGCAGGCAGAGACTGACAA | CAGCAGCAGCAGCAGCAGAG |
| *GALNT6* | GGGAGGGTTCAGGGCAGCAATG | GGGAGGGTGGGCGTAGAGATGG |
| *GALNT7* | GGCTAGTGGTCCTCTGGTCTTCC | CTCTGTCTTCCCTCATCCTGCTCA |
| *GALNT8* | CTCCAGACCTGTAGCACGCAAGT | TGATGACCCAGATCCAGCACCAT |
| *GALNT9* | GGCAGACACAGCAGGACA | TGTTTGAGTTGGCATCACTGTC |
| *GALNT10* | AAGGAGCCAGCCAGGTGTAATAC | CAGTTGTGATGATGGTGGGAGGTT |
| *GALNT11* | GCCACGGGTCAGGAGGAT | TGAAGAGGAGCCATCGCAGAT |
| *GALNT12* | CAGGCTGCGAGGAAGGAGTC | GGTGGGTTCTGGTCACTGCTTAG |
| *GALNT13* | CGTCCACACCACTCACCACACAT | TGGGGCAGAGGGAACAAAACACT |
| *GALNT14* | TCGTCGTCAACCCATGTGAGTCCT | GCTGCCCAGTTTCCAGTCTGTTCT |
| *GALNT15* | AACACTGGACTTGGGCTCTG | CACCTGCTCCTGCCTGAC |
| *GALNT16* | CCTGAACCTCTGCTCTGGATTG | TGCGGACCACAGACACTTG |
| *GALNT17* | AACAGCGGAGCGTGAGAG | GGAGTCCAGCCAGGAAGTG |
| *GALNT18* | CACCGTGGATGATGATGACAAC | AACTCCAGGTCGCTATTCTCC |
| *GALNT19* | AGACGCCTTCCACGAGAT | CACGATGTCCTCCAGCAG |
| *GALNT20* | CCTCCTGTGTCTCATCTCTTGC | TGGTCCTCACTGCCTCCTT |
